# Automatic Synthesis of Anthropomorphic Pulmonary CT Phantoms

**DOI:** 10.1101/022871

**Authors:** Daniel Jimenez-Carretero, Raul San Jose Estepar, Mario Diaz Cacio, Maria J Ledesma-Carbayo

## Abstract

The great density and structural complexity of pulmonary vessels and airways impose limitations on the generation of accurate reference standards, which are critical in training and in the validation of image processing methods for features such as pulmonary vessel segmentation or artery–vein (AV) separations. The design of synthetic computed tomography (CT) images of the lung could overcome these difficulties by providing a database of pseudorealistic cases in a constrained and controlled scenario where each part of the image is differentiated unequivocally. This work demonstrates a complete framework to generate computational anthropomorphic CT phantoms of the human lung automatically. Starting from biological and image-based knowledge about the topology and relationships between structures, the system is able to generate synthetic pulmonary arteries, veins, and airways using iterative growth methods that can be merged into a final simulated lung with realistic features. Visual examination and quantitative measurements of intensity distributions, dispersion of structures and relationships between pulmonary air and blood flow systems show good correspondence between real and synthetic lungs.

## 1. Introduction

During the last few decades, there has been increasing interest in the design of synthetic images, and especially those comprising anthropomorphic reference standards, to support the development of new image processing algorithms by training and evaluating them in a constrained, simplified, and controlled scenario. However, the extraordinary complexity of pulmonary structures has been an impediment to obtain sufficiently realistic synthetic images of the lungs. Phantom images could be useful in the development of methods for processing internal structures (vessels, airways, and lung parenchyma), but also for any other approaches, such as the design of registration algorithms that could deal with changes in the lung and internal structures.

The great density and structural complexity of pulmonary vessels and airways impose limitations on the generation of accurate reference standards for the evaluation of image processing methods specifically designed for these structures. Currently, schemes for validating segmentation algorithms are usually constrained to a limited number of points with no connectivity or size information [1], which results in a relatively limited evaluation that could be improved by using information about the whole tree structure. Moreover, the design of pulmonary AV segmentation/separation algorithms, considered as one of the main future challenges [2], is being delayed by the difficulty in training and/or validation of the methods because of the absence of properly labeled images in which both trees are differentiated correctly.

Because manual segmentation and annotation of vessels and airways are extremely time-consuming tasks, the use of a tool to generate realistic pulmonary phantoms would overcome the difficulties by providing a database of realistic CT cases and a corresponding gold standard. Moreover, this simulation framework would permit synthesis and posterior evaluation over a wide range of conditions (levels of noise, artifacts, pixel size, contrast, and the inclusion of pathologies).

The most important and challenging component in simulating lung imaging is the generation of synthetic vasculature and airways. Simple computational vascular phantoms include “ad hoc” images specifically designed for testing algorithms under very simplified and constrained conditions. Typically, they are tubular-like structures with low levels of complexity and realism such as the ones presented in [3]. Complex computational vascular phantoms have been addressed using two main different schemes: a) Lindenmayer systems, based on grammatical rules that construct complex structures in an accumulative manner [4]; and b) iterative growth methods based on perfusion models and growing vascular structures that connect new selected terminal nodes with the principal structure [5, 6, 7, 8, 9]. These approaches have always been based on some vascular network model that takes into account structural (size, location) and functional (blood flow) features following physical hemodynamic laws [10, 11]. Iterative growth methods are more extended because they are easy to connect with hemodynamics, control the volume of vasculature, and constrain the generation of the tree structure. Specific computational airway models have been developed using similar strategies based on fluid dynamics [12, 13, 14].

To our knowledge, no proper computational CT phantoms of the whole lung have yet been created. Only some approximations based on deformations from original CT images [15] or labeled images obtained using different injected contrast materials during acquisition [16] have been presented. Additionally, [17] described a method to generate and include supernumerary vessels into a manually labeled AV segmentation image. This strategy was also used to complete the pruned airway tree extracted from real CT images [18]. Different kinds of approaches have presented complex functional models of the whole lung, and vessel and airway structures [19, 20]. However, these works suffer from at least one of the following limitations: they do not address the generation of a complete image phantom with the specific characteristics of a certain imaging modality; they do not differentiate all internal structures (vessels, airways, and parenchyma); and/or make impossible the systematic generation of multiple and diverse phantoms to create a standard reference dataset.

Here, we present a framework based on VascuSynth [9] to generate anthropomorphic pulmonary CT phantoms of the human lung automatically. The system is able to create synthetic vessels (arteries and veins independently), representing the pulmonary circulation, and airway trees based on prior knowledge that characterizes anatomical and topological relationships of pulmonary structures as well as their corresponding image features, which will be merged into a final simulated lung with realistic features.

## 2. Methods

Brigham and Women’s Hospital obtained approval from the Partners Human Research Committee and all subjects provided written informed consent, allowing the use of CT data in this study.

The generation of synthetic pulmonary CT images is detailed in the following two sections. First, we present the generation of simplified pulmonary phantoms by simulating bronchopulmonary segments. Second, we extend the idea to create more complex synthetic images: the final anthropomorphic pulmonary CT phantoms. Fig. 1 shows a scheme of the whole generation process, where the synthesis of anthropomorphic pulmonary phantoms includes an additional previous step over the general workflow used for the generation of bronchopulmonary segment phantoms.

**Figure 1:**
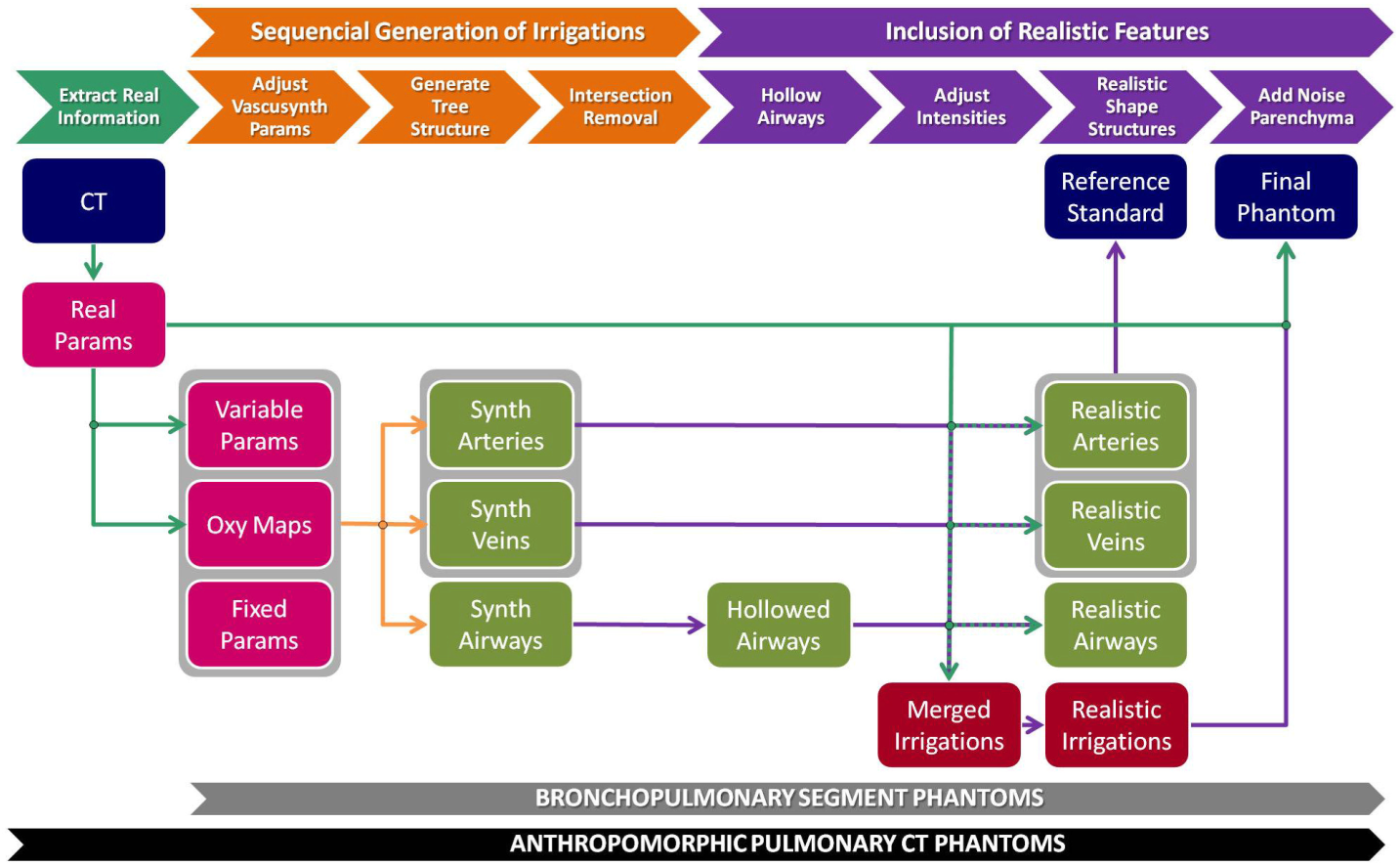
Pipeline of pulmonary phantom generation. The first stage (orange) includes processes to generate the arterial, venous, and airway tree pathways. The second stage (purple) comprises different steps to provide realistic features to the final phantom. An additional previous stage (cyan) is introduced when generating anthropomorphic pulmonary CT phantoms, but not in bronchopulmonary segment phantoms, to extract information from the real CT image that will be used to adapt some of the parameters. Blue boxes represent input and output images; pink boxes represent input objects and parameters; green boxes are intermediate images for a single structure; and red boxes represent images with the three subtrees corresponding to the pulmonary air and blood flow systems together.

### 2.1. Bronchopulmonary Segment Phantoms

The first step to generate synthetic images was the development of phantoms with simplified pulmonary-like structures. To that end, we focused on the repetitive anatomical and functional structure in the lungs that exhibits a vascular pattern: the bronchopulmonary segment.

Arteries to a bronchopulmonary segment are distributed within the segment along with the branches of the ventilating bronchus, whereas veins lie in the intervals between segments [21]. The idea behind the generation of a bronchopulmonary segment phantom is the synthesis of tubular structures that represent pulmonary arteries, veins, and airways following this anatomical pattern. However, one of the most important features to pursue in the generation of phantoms is the interest in creating multiple and different synthetic images from the same initial conditions. Therefore, there should be a balance between the randomness of the algorithm and the relationship between the air and blood flow systems, while maintaining both principles during generation.

We can outline the generation of bronchopulmonary segment phantoms in two main stages: a) generation of the pulmonary air and blood flow systems; and b) inclusion of realistic features.

#### 2.1.1. Sequential generation of arterial, bronchial, and venous flow systems

Initially, the three flow systems are generated as tubular-like structures using the same strategy. VascuSynth [9, 22] is an open source software to synthesize three-dimensional (3D) tubular tree structures iteratively based on a physiological model of flow conservation; it was therefore a suitable starting point for our purpose. Because VascuSynth was developed for the simulation of divergent flows and for creating opaque tubular structures, it can only be used to generate the pulmonary arterial system directly. However, we can simplify the representation and model the venous system based on the arterial one by considering that its direction of flow is actually the opposite. For simulating the airway system, it can be reduced to an arterial one ensuring the appropriate emptiness of the structure. Therefore, by selecting the proper parameters, each flow system can be produced independently using VascuSynth. Nevertheless, some restrictions and constraints need to be applied to comply with the desired vascular pattern.

Construction of the phantom is performed sequentially by taking into account the anatomical relationships between flow systems generated in the previous steps. Given a specific oxygenation map, the arterial tree is generated first within the area of the simulated segment, locating it in the central part preferentially; second, airways are generated close and parallel to arteries; finally, veins occupy the external parts of the segment, relatively far from arteries and airways. For each structure, the process can be reduced to two steps: adjusting VascuSynth parameters and generating a final image of the tree structure by running VascuSynth software (Fig. 2). The following subsections describe this procedure in detail.

**Figure 2:**
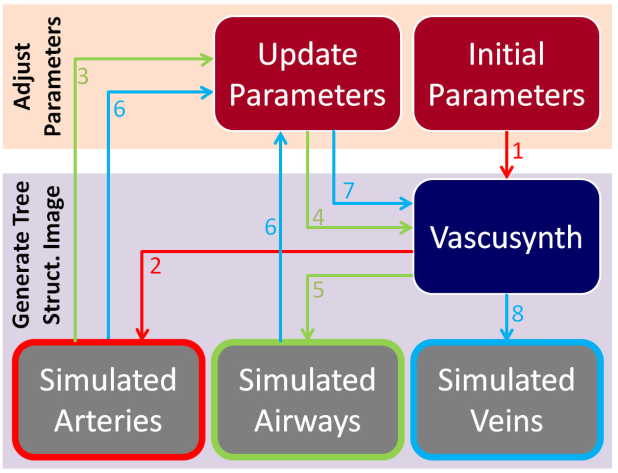
Workflow for the generation of a synthetic bronchopulmonary structure for arteries (red arrows), airways (green), and veins (blue). Numbers indicate the order in the workflow.

##### Adjust VascuSynth Parameters

VascuSynth creates tubular-tree structures starting from some user-defined physical parameters that determine the tree hierarchy, branch properties, and bifurcations [9].

- Fixed Parameters: related to physical and physiological variables of the pulmonary system and the blood. Initial and final pressures (*P*_*ini*_ = 25 *mmHg* and *P*_*fin*_ = 10 *mmHg*, respectively) and viscosity of the fluid (*η* = 36 *mP a · s*) were based on real reported values [23]. Initial flow (*Q*_*flow*_ = 138.83 *cc/min/g*), initial radii parameter (*λ* = 2), radii factor (*γ* = 2.55), a parameter controlling the length of branches (*μ* = 2), the length factor of branches (*D*_*T*_ = 1 *mm*), and the number of closest branches to consider during construction of the bifurcation (*k* = 5) were fixed empirically taking into account a priori biological information.
- Variable Parameters: several parameters change depending on the structure to be simulated or the complexity of the output image we want to achieve. For construction of the bronchopulmonary segment phantoms, the following parameters were fixed: the perfusion point (*p*_0_) determines the location where the whole tree structure will start to develop 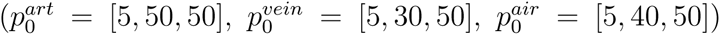; the size of the image (*S* = 101 *×* 101 *×* 101) and the number of terminal points (*N*_*f*_) establish the structural complexity. In our experiments, arteries and veins share the same number of terminal points 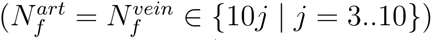 and airways have a percentage of AV terminal points (determined by the factor of airways, *F A* ∈ {1, 2, 3}, 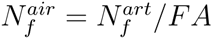), when trying to simulate the common circumstances in CT imaging where the number of visible airway generations is always less than the number of vessels. A seed number (*R*_*seed*_) was selected pseudorandomly in every simulation to introduce the aleatory component of the system.
- Oxygenation Maps: the most important parameters to achieve our objective are the nutrient maps: an oxygenation map (specifying the demand for oxygen at each point of the 3D space) and supply map (determining the effective values of the oxygenation demand at each point). Default values for the supply map were used, so the oxygenation map acts as a probability map for tree growth. It is represented as a 3D scalar volume that covers the entire lung parenchyma where perfusion exists, with values from 0 (no demand) to 1 (maximum demand) for oxygen. During each iteration in the construction of the tree, a candidate terminal node is chosen according to the oxygen demand map, creating a bifurcation from an existing branch to supply this terminal node. Ultimately, the probability of a specific point to be a terminal point depends on its oxygen demand value and the existence of an oxygenation path connecting this candidate terminal point with the subtree synthesized previously [9]. Since the initial whole volume to be synthesized represents just a single bronchopulmonary segment, arteries should be distributed within the central part of the volume, accompanying the airways. Contrarily, veins should occupy the external parts of the volume. Thus, the oxygenation maps for arteries (Fig 3a) and airways present with higher values of oxygenation demand (and therefore show a higher probability of developing branches) in the central area, and lower values at the segment limits. The opposite strategy is followed to construct the oxygenation map for veins (Fig. 3c). To include some uncertainty in the tree generation and smooth the oxygenation map, intermediate values of oxygenation demand are made to occupy middle regions by applying a gradient-like distribution using chessboard distances. Additionally, arteries and airways should comply with the parallelism and proximity restrictions present in the bronchopulmonary segment structure. To obtain these, the oxygenation map for airways (Fig. 3b) must be modified to promote airway growth around the arteries. Mathematical morphology functions and distance transforms create maximum values of oxygen demand in areas located at a distance 2*r*_*max*_ from the previously generated arteries, becoming gradually lower until a maximum distance of *r*_*max*_ is reached. Finally, because two different structures should not be located at the same position, a null oxygenation demand is established in areas where previously generated structures are placed (plus a security margin) to minimize overlapping. Fig. 3 shows the final oxygenation maps and the corresponding tubular structures generated from them.

**Figure 3:**
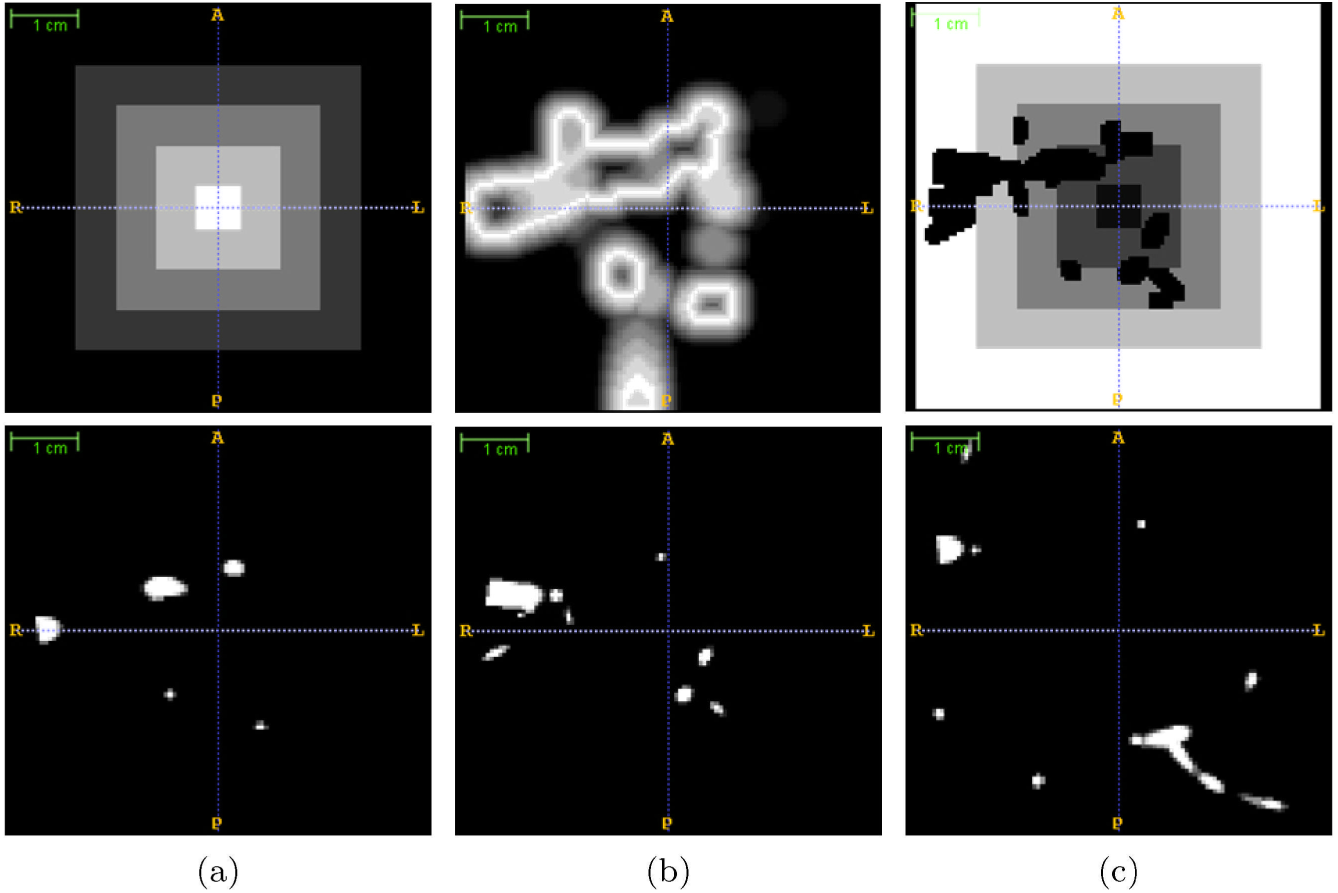
Example of oxygenation maps used in the synthesis of a bronchopulmonary segment phantom. The first row displays the central slices of the 3D oxygenation maps for arteries (a), airways (b), and veins (c); bright/dark areas indicate high/low levels of oxygenation demand, and therefore, the probability of growing structures there. The second row shows the 3D structures generated from the corresponding oxygenation maps, which are used to define subsequent oxygenation maps for the following structures.

##### Generate Tree Structure Image

By following the workflow presented in Fig. 2, one volumetric image for each tubular structure is generated by running VascuSynth with specific defined parameters and oxygenation maps (Fig. 3).

##### Intersection Removal

Although this construction of the three flow systems tries to avoid overlapping of structures, the way in which VascuSynth performs could cause partial superpositions in some areas that need to be corrected by distributing joint voxels between the structures and by removing disconnected branches that could appear after the process. Thus, the output could display touching structures that, far from being undesirable, simulate the intertwining nature of vessels and airway trees present in real cases.

#### 2.1.2. Inclusion of realistic features

To construct the final phantom, some postprocessing methods need to be applied to create a more realistic CT simulation of the bronchopulmonary segment, including the hollowing of airways, the deformation of structures, and the inclusion of parenchyma and noise.

##### Hollowing of Airways

Unlike vessels, airways are hollow structures. Therefore, we need to transform the opaque tubular structures obtained with VascuSynth into empty structures. Airways share the following premises.

- Intensity values of airway lumens are similar to the lung parenchyma in CT images, but partial volume effects increase the differences in *HU*. Airway walls and vessels (with no contrast media) display similar intensity values [24].
- The thickness of airway walls is variable and decreases with the size of the airway. Studies such us [25] assessed the ratio between luminal diameter and wall thickness (*LD/W T*) in different generations of an airway tree. From these values, we can compute the ratio between the wall thickness and the diameter of the airway for each generation (*WT/*(*LD* + 2 · *WT*) = *WT/D*).
- The small scale of airway walls produces partial volume effects and, consequently, a reduction in actual intensity values at the wall voxels.
- These three features mean that the bifurcation levels of airway trees in CT images become indistinguishable from lung parenchyma earlier than do vessels.

The first point is addressed in the next subsection, where intensity values are adapted to realistic ones. The last point was addressed previously by using a variable *FA* parameter with a value greater than 1 that decreases the density of airways with respect to vessels. The second and third points are addressed in this section.

Given that VascuSynth generates tree structures by creating straight segments that connect initial, bifurcation, and terminal points with a constant radius along each segment, we can hollow the airway structure by using a 3D erosion subtraction strategy in each segment of the tree, determined by its *WT* value, obtained from the caliber of the segment and the *WT/D* ratio. Because *WT* probably does not correspond to an exact integer in terms of voxels, the process introduces a simulation of partial volume effects.

##### Adjusting Intensities

Because the original structures obtained with VascuSynth are represented by grayscale images in a realistic way (brighter in the centers and gradually darker toward the walls), a simple linear transformation of *HU* is applied to obtain realistic values found in pulmonary vasculature (and airway walls) in CT images of human lungs (35 *–* 45 *HU*) [26].

##### Realistic Curvature of Vessel and Airway Trees

To incorporate a higher degree of realism, we can bend the tubular segments, adding curvature to the structures. We use a deformation algorithm based on nonrigid B-spline transformation [27, 28] to deform vessels and airways slightly. The basic idea is to create a nonrigid transformation where initial, final, and bifurcation points from the three flow systems remain unaltered, and the segments of the trees are deformed by moving their middle points with a pseudorandom displacement vector with a maximum magnitude of *d*_*max*_. The generated B-spline grid is regularized by making it diffeomorphic (smooth and invertible). Fig. 4 shows an example of the deformed version of a subtree.

**Figure 4:**
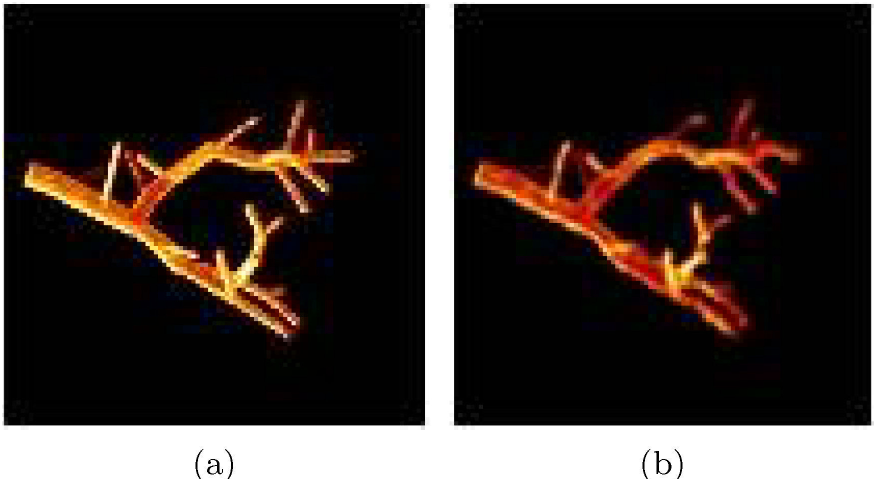
Example of the application of a deformation algorithm to a tubular tree structure (a). The resulting image is shown in (b) using a value of *d*_*max*_ = 3.

##### Final Reference Standard

These three independent images corresponding to the vascular and airway trees can now be merged into a single labeled image that will serve as a reference standard for the final synthetic CT image of this bronchopulmonary segment.

##### Adding Parenchyma and Noise

After merging the three structures with adjusted intensity values, we need to include a realistic background that will correspond to lung parenchyma. To simulate a pseudorealistic texture and intensity values of lung background (associated with the inclusion of tiny structures and the variable densities of blood and air), we assign random values of 800 *±* 150 *HU* to voxels that do not correspond to any of the subtrees. Then, a Gaussian blurring with standard deviation *σ* = 1.5 *mm* simulates a reconstruction kernel, smoothing and adding texture to the synthetic parenchyma.

Finally, some characteristic noise sources are added to the image (Gaussian white noise of standard deviation *σ* = 20 and small-density salt-and-pepper noise) to create the final bronchopulmonary phantom.

### 2.2. Anthropomorphic Pulmonary CT Phantoms

Although bronchopulmonary segment phantoms follow the theoretical structure of real segments in the lung, they represent only a certain region of the whole lung. Moreover, they assume an ideal situation in which airways run parallel to arteries covering the whole area of the segment. However, this is not a fully realistic situation in CT images, where airways become indistinguishable from lung parenchyma earlier than do vessels when they become smaller (more noticeable in peripheral areas of the lung). It is true that the *FA* factor tries to deal with this problem, but the effect is just a reduction of the density of the airways with respect to vessels, and does not take spatial restriction into account.

To create the final pulmonary CT phantoms in the most realistic way, we have used a set of real labeled CT cases. From these CT images and their arterial, venous, and airway tree segmentations, we can extract information properly to adapt some parameters from the previously explained workflow (Fig. 1); that is, adjust the VascuSynth parameters and adjust the intensities of air and blood flow systems and the parenchyma. Following this procedure and taking into account the pseudorandom nature of the simulation, the method is able to create multiple phantoms that fit particular features of a certain real case.

#### 2.2.1. Noncontrast Pulmonary CT Data

The three CT cases used in this work were acquired as part of the COPDGene study [29] from Brigham and Women’s Hospital (BWH) in Boston, and obtained through dbGap (http://www.ncbi.nlm.nih.gov/projects/gap/cgi-bin/study.cgi?study_id=phs000179.v4.p2). The data were anonymized by the COPGene study data coordinating center before sharing them. Apart from the noncontrast CT images, segmentations of lungs [30], vessels [31], and airways [32] were available for these three patients suffering from chronic obstructive pulmonary disease (COPD), resulting in six different and completely segmented lung cases. A manual separation and labeling of arterial and venous structures from vessel segmentations were carried out by two trained experts using ITK-Snap (www.itksnap.org).

#### 2.2.2. Extraction of Information from Labeled Pulmonary CT Images

Given the lung, vessel, and airway segmentations, some structural and image-based features were computed in the proper places.

- Location, size, and lung shape from the lung segmentation image.
- Intensity distributions. Mean intensity and standard deviation in arterial (*μ*_*art*_, *σ*_*art*_), venous (*μ*_*vein*_, *σ*_*vein*_), and parenchymal (*μ*_*lung*_, *σ*_*lung*_) locations. Because of the difficulty in getting accurate locations of airway walls, *μ*_*air*_ and *σ*_*air*_ are fixed to the experimental values used previously in the generation of bronchopulmonary segment phantoms.
- Terminal points in arteries, veins, and airways. A skeletonization [33] of the segmented structure is applied, and the number of terminal points is obtained by detecting voxels with only one neighbor belonging to the skeleton.
- Root location. Given the terminal points, the root corresponds to the branch with larger diameter. Sphere inflations are used to detect this location.

#### 2.2.3. Adjust VascuSynth Parameters

VascuSynth simulation is adapted following the information extracted from the real CT case. Global and structure-dependent parameters of VascuSynth are adjusted correctly: size of the image (*S*), pixel spacing (*ps*), number of terminal points (*N*_*f*_), perfusion or initial points (*p*_0_), and oxygenation maps (*O*_*map*_).

The first three parameters take values extracted from the image as already described. Perfusion points (*p*_0_) are fixed to locations with high values of oxygenation demand and very close to the root coordinates extracted in the previous subsection. Oxygenation maps are created exploiting the information from the labeled segmentations of the flow systems.

##### Oxygenation Maps

Following the bronchopulmonary approach, anthropomorphic oxygenation maps retain the use of prior information about spatial relations between air and blood flow systems. However, some modifications are introduced to generate more realistic oxygenation demands at the complete lung level (Fig. 5).

**Figure 5:**
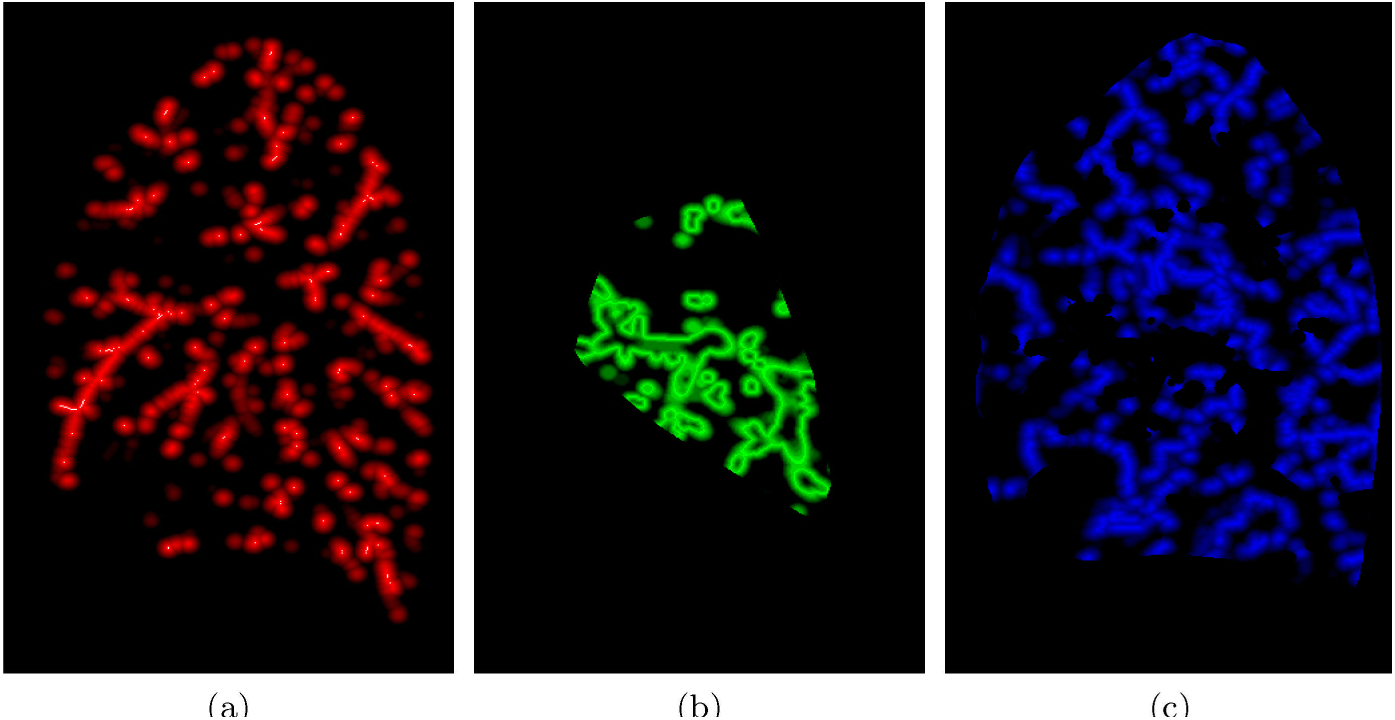
Oxygenation maps in anthropomorphic pulmonary phantoms. Sagittal views of arterial (a), bronchial (b), and venous (c) oxygenation maps.

- Arterial Oxygenation Map: This oxygenation map only depends on the real spatial distribution of manually segmented arteries. The basic idea is to create a smooth probability map inside the lung with high values close to locations of real arteries, and small values far from them. A distance transformation from the center line of the segmented arteries is applied and normalized to obtain a maximum value (1) in each of the center lines and a null value farther than a user-defined maximum distance (*dmax* fixed to 10 *mm* in our experiments).
- Airway Oxygenation Map: The method is the same as the one used for the bronchopulmonary segment phantoms. However, to include the restriction of airways becoming indistinguishable from lung parenchyma earlier than do vessels when they become smaller, we constrain the generation of the airway tree in the convex hull of the real bronchial tree segmentation, extracted from the real CT image, by assigning a null oxygenation demand in the areas outside this region.
- Venous Oxygenation Map: The desired spatial structure of veins is as far as possible from arteries and airways, so its oxygenation map is essentially the inverse of the arterial/bronchial one. To create it, skeletonization, distance transformation, and normalization are applied to obtain a maximum value in the points located farther from the previously generated arterial and bronchial systems (desired venous center lines) and a minimum value (*ox*_*min*_=0.001) in points located farther than *dmax* from that center line. A null oxygenation demand is fixed in areas outside the lung and places already occupied by the synthetic arteries and airways previously created.

#### 2.2.4. Adjusting Intensities

Instead of applying common *HU* values found in real CT images, intensity information extracted from the specific real case is used to generate similar distributions in its associated final phantoms. Thus, synthetic arteries, veins, airway walls, and parenchyma would fit, respectively, a distribution of mean *μ*_*art*_, *μ*_*vein*_, *μ*_*air*_, *μ*_*lung*_ and standard deviation *σ*_*art*_, *σ*_*vein*_, *σ*_*air*_, *σ*_*lung*_.

#### 2.2.5. Inclusion of Lung Phantom into the Real CT

Finally, to construct a complete CT image, the lung region from the real CT is substituted with the generated final pulmonary phantom. In that way, the final image maintains the global features from the original one, but with a different structure of interest.

## 3. Results and Evaluation

### 3.1. Bronchopulmonary Segment Phantoms

A database of different bronchopulmonary segment phantoms was created by changing the initial variable parameters with the values shown in the corresponding section. Four cases for each combination of *N*_*f*_ and *FA* parameters were generated resulting in a total of 96 bronchopulmonary segment phantoms. Fig. 6 displays the difference in airway density obtained by changing the *FA* parameter, and Fig. 7 displays different cases obtained by varying the number of terminal points *N*_*f*_.

**Figure 6:**
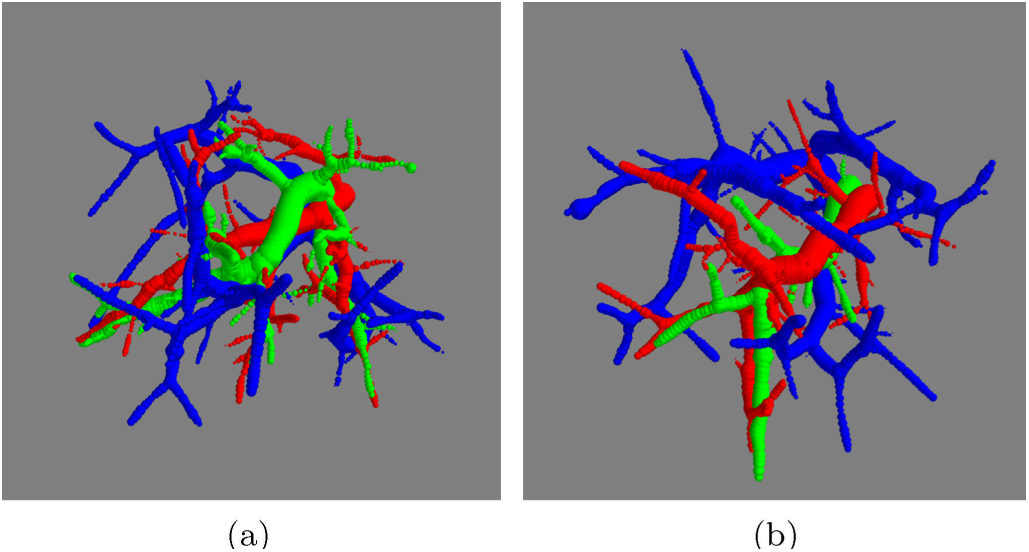
Effect of randomness and *FA* factor in bronchopulmonary segment phantoms, displaying arteries in red, veins in blue, and airways in green, with *N*_*f*_ = 50. (a) *FA* = 1 *⇒ N*_*f*_ (*air*) = 50, (b) *FA* = 3 *⇒ N*_*f*_ (*air*) = 17. Although both examples share the same parameters for the synthesis of arteries, the generated arterial flow systems have different structures because of the change in random seed *R*_*seed*_.

**Figure 7:**
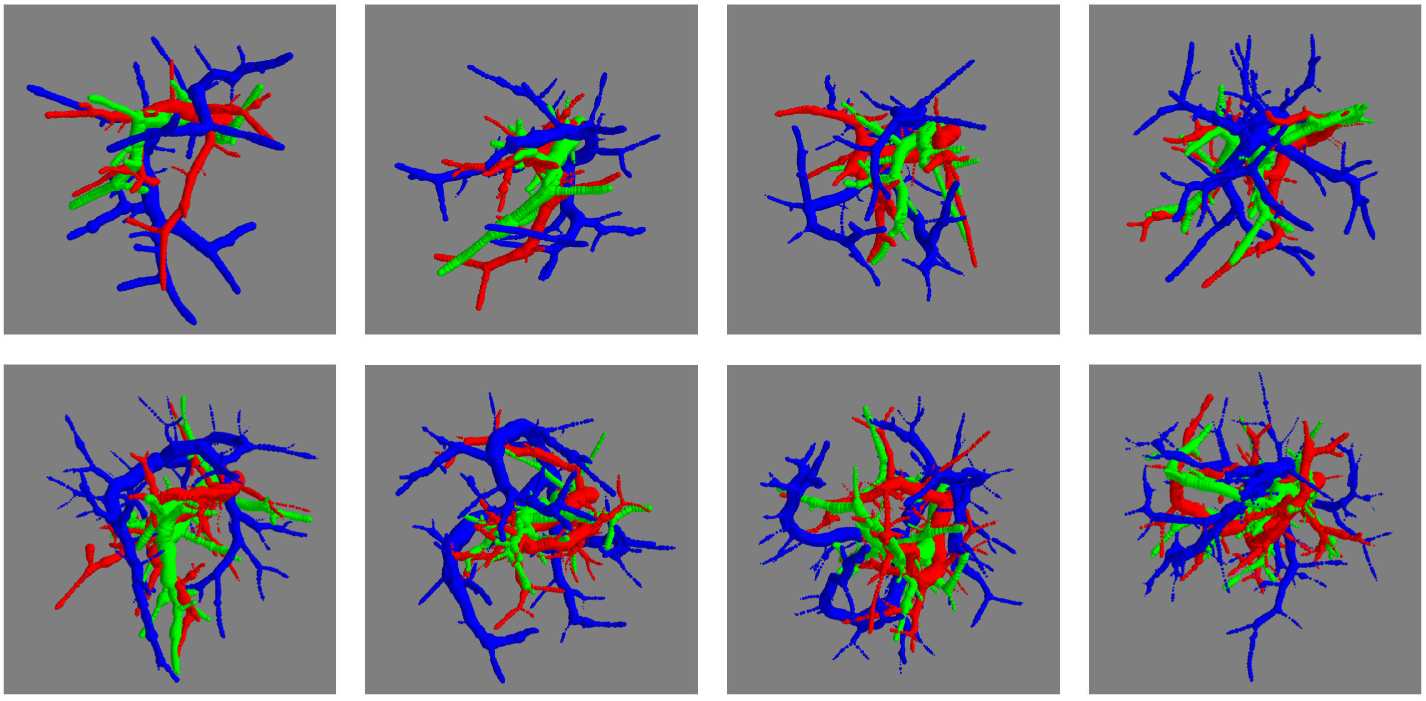
Effect of the numbers of terminal nodes in bronchopulmonary segment phantoms. From left to right, up to down, arterial (red), venous (blue), and airway (green) structures with *N*_*f*_ = {30, 40, 50, 60, 70, 80, 90, 100} and *FA* = 3.

### 3.2. Anthropomorphic Pulmonary CT Phantoms

Four phantoms were generated from each real lung of the three noncontrast CT cases available. The different values of the random seed guarantee diversity in the four cases sharing the same inputs. Therefore, *M* = 24 final anthropomorphic pulmonary phantoms were generated. Fig. 8 shows different final synthetic CT images obtained from the same real CT case and Fig. 9 displays 3D reconstructions of the pulmonary structures generated synthetically from different cases. Supplementary video (S1 Video) shows a slice-by-slice fly-through of a resulting phantom CT image as well as the image with the overlay of the inner structures.

**Figure 8:**
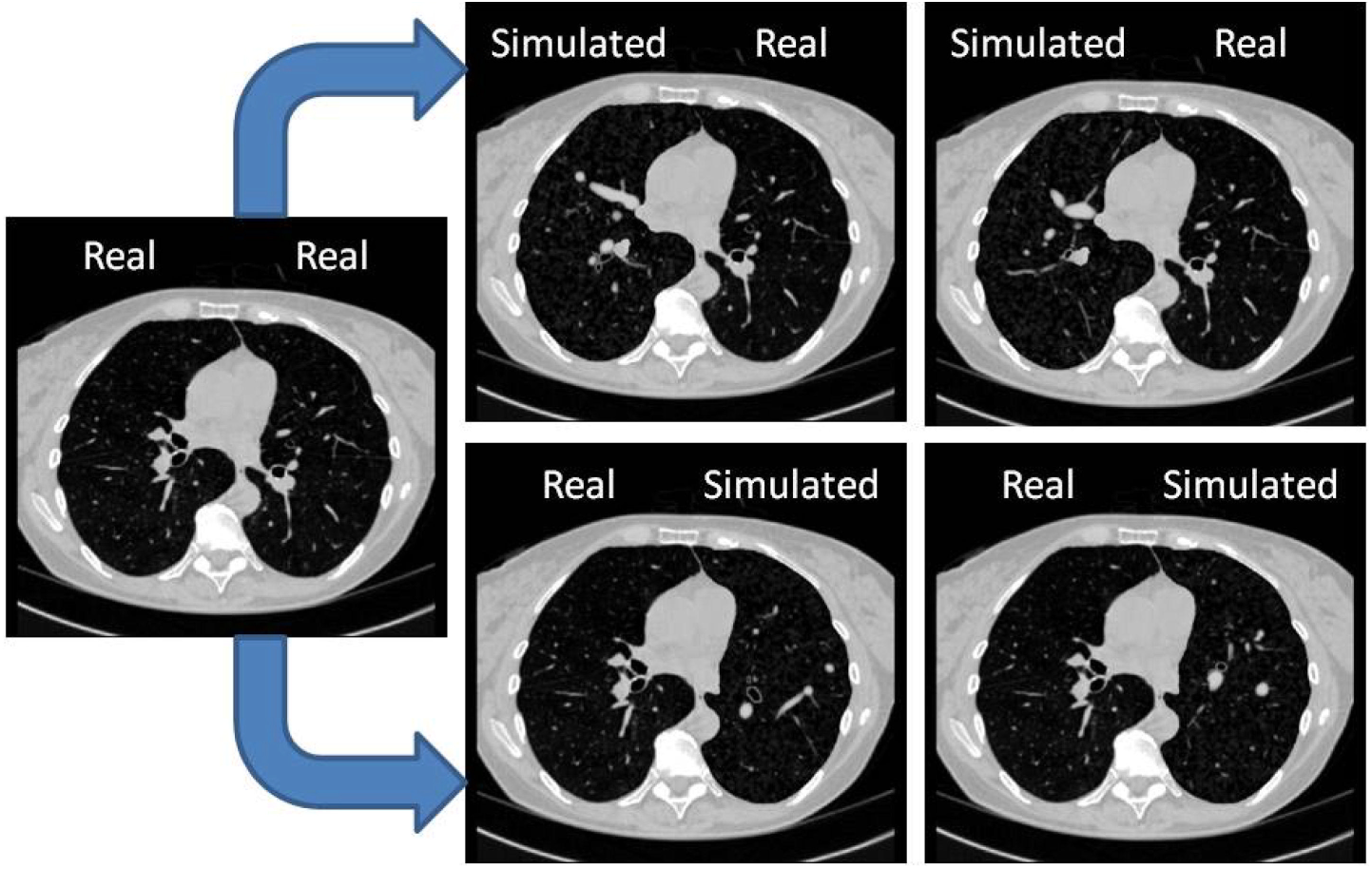
Results of anthropomorphic final phantoms, showing two CT images with synthetic right lungs (top row) and left lung (bottom row) obtained from a real CT case (left). L=-400, W=1500 *HU*.

**Figure 9:**
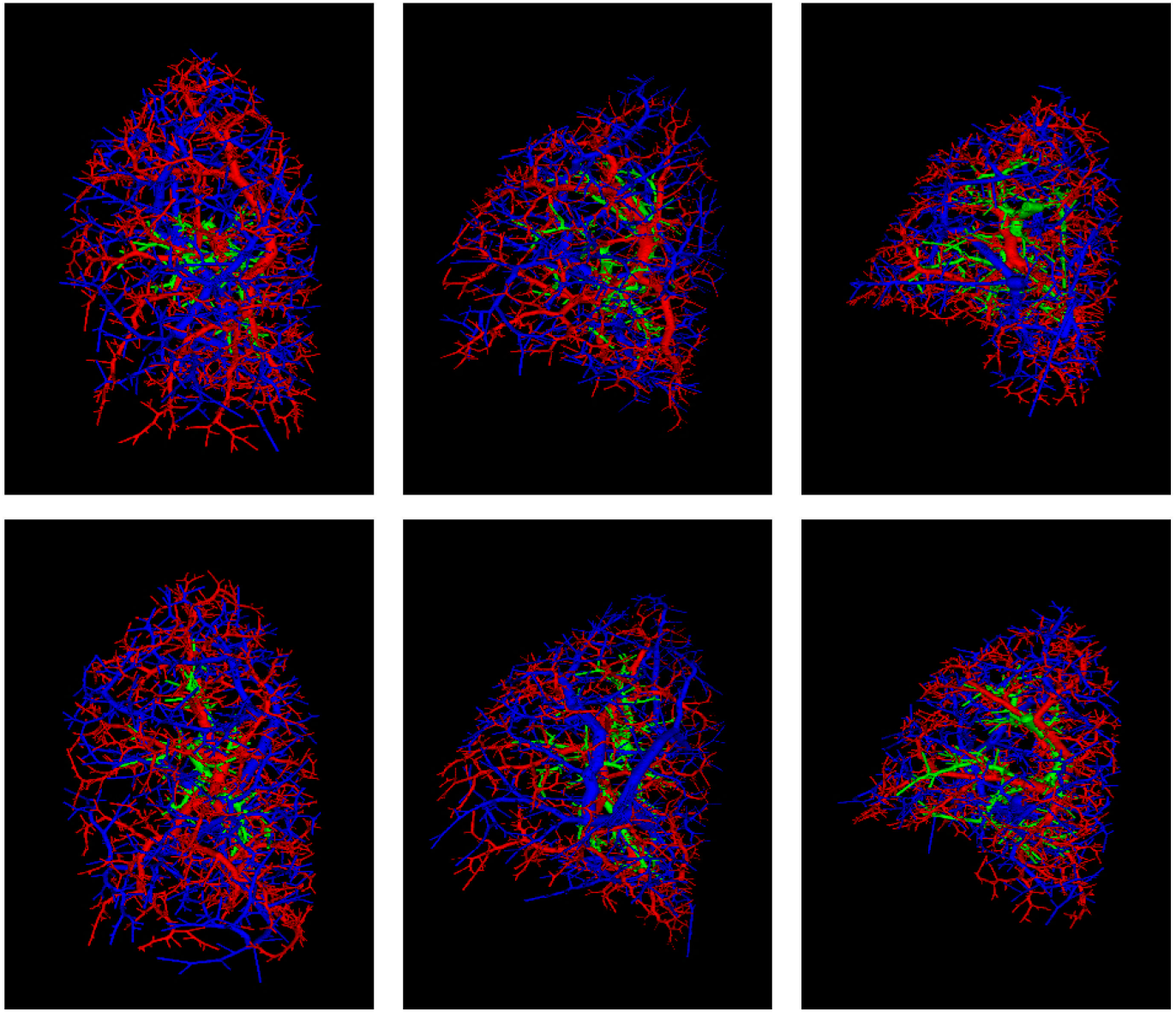
Synthetic arteries (red), veins (blue), and airways (green) conforming the reference standard for the anthropomorphic pulmonary CT phantoms. The first column corresponds to synthetic air and blood flow systems of two phantoms generated from the left lung of one real case. Similarly, the second and third columns correspond to phantoms from the left and right lungs, respectively, from another real case.

#### 3.2.1. Validation Strategy

A visual examination of the synthetic pulmonary phantoms demonstrates a good similarity of the simulated lungs to the real ones (Fig. 8). Nevertheless, it is just a qualitative comparison. Some quantitative measurements could provide more objective information to evaluate the generated phantoms.

Different evaluation measures were extracted from real and synthetic images. Intraclass variability in real cases (real vs real) and interclass variability (real vs phantom) were computed using pairwise similarity metrics. A comparison between them allows us to assess whether the produced synthetic phantoms are within the range of variability of real lungs.

##### Evaluation Measures

Because of the typical anatomical variability present in pulmonary vessels and airways, direct voxel-based comparisons or measurements taking into account spatial locations of structures are not suitable. For that reason, the evaluation measures are based on global and local features of the image using intensity and distance histograms as estimates of the corresponding probability density functions (PDFs).

- Intensity distributions: PDFs of *HU* within the lung (Fig. 10a).
- Dispersion of structures: the density and sparsity of internal structures can be estimated measuring the size of “holes” between internal structures by computing the PDF values of euclidean distances from parenchymal locations to vessels/airways (Fig. 10b).
- Relationship between arteries and airways: PDF values of euclidean distances from arterial points to airway locations (Fig. 10c). Although this spatial relation information was used initially in the generation of phantoms, it is just a theoretical tendency. The variability introduced by the random component and posterior deformation transformation could bias the general tendency in these synthetic images.

**Figure 10:**
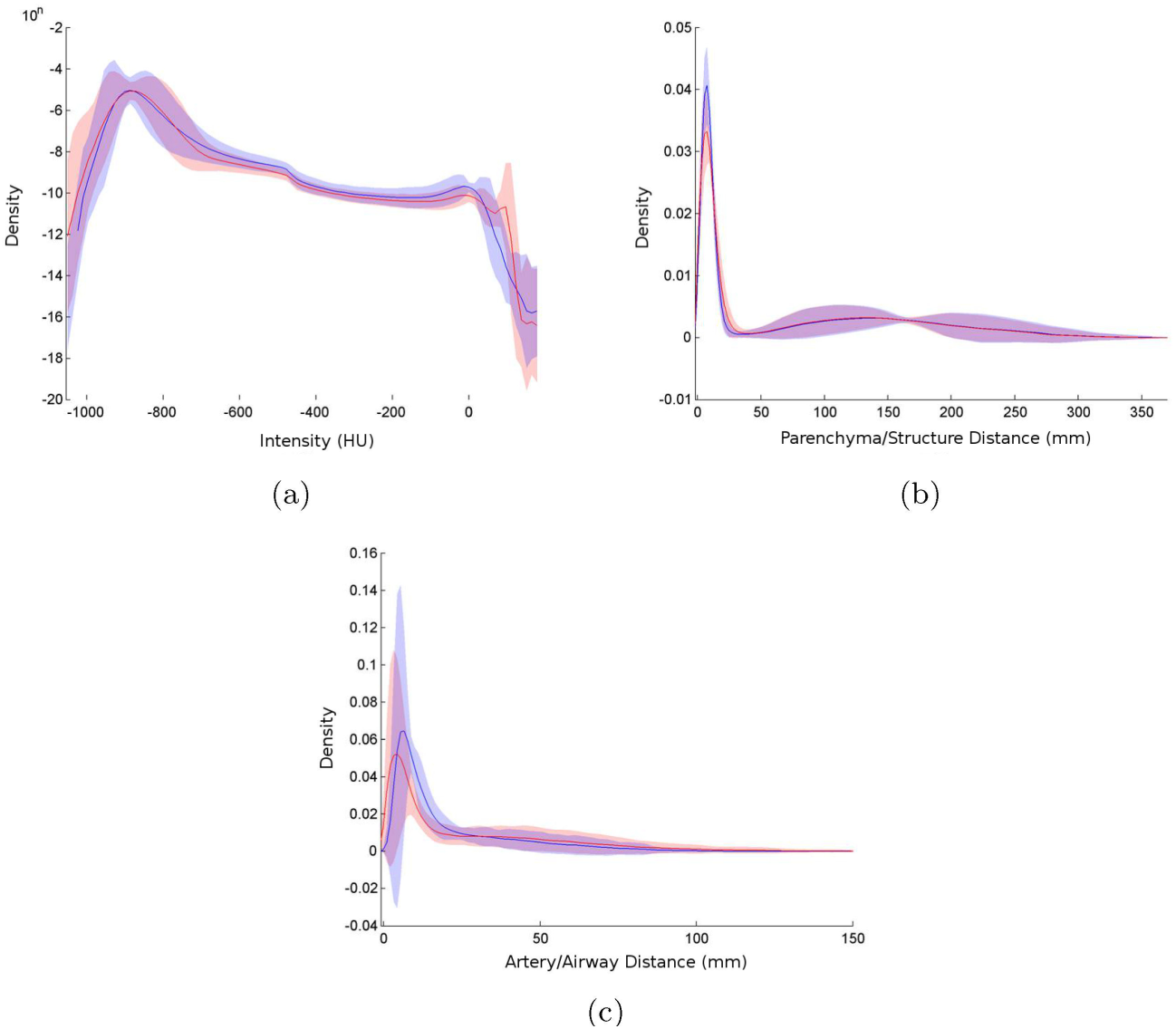
Histograms of evaluation measures showing mean probability density functions (lines) with 95% confidence intervals (shadows) for real (blue) and synthetic (red) lungs. (a) Intensity distribution. (b) Dispersion of structures. (c) Relation between arteries and airways.

##### Similarity Metrics

The similarity/difference between two density functions was estimated using histogram-based match distance and Kolmogorov–Smirnov distance [34]. Pairwise comparisons were performed to compute intraclass distances in real cases (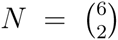) and interclass distances (real vs phantom, *N* = 6 × 24). Mean and standard deviation of the intraclass and interclass pairwise distances were reported for each evaluation measure.

##### Statistical Analysis

The Wilcoxon rank-sum test was used to compare intraclass variability in real cases and interclass variability, complemented with Cohen’s *d* [35] to measure the effect size and area under the ROC curve (AUC) to measure the degree of separability between populations.

### 3.2.2. Validation Results

The statistical analysis (Table 1) reports a statistically nonsignificant difference (*p >* 0.05 and fairly small values of Cohen’s *d* and *AUC*) when comparing real vs real and real vs phantom similarity measures. This means that interclass differences are similar to those found between real cases; therefore, synthetic phantoms are within the range of variability of real lungs for all three evaluation measures used for this comparison.

**Table 1:**
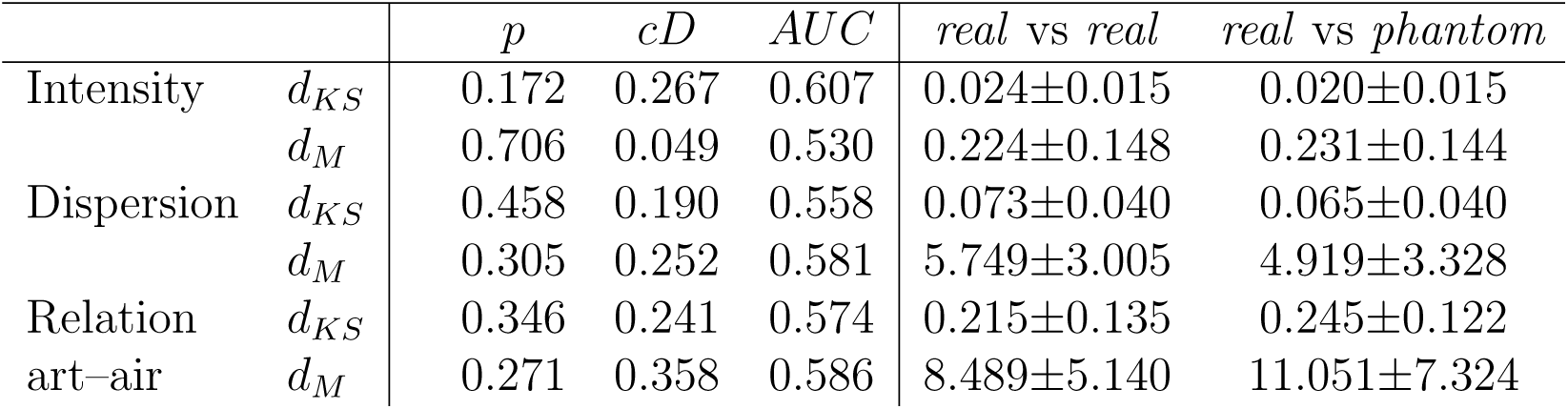
Statistics comparing intraclass similarity measures in real cases (real vs *real*), and interclass similarity measures (*real* vs *phantom*). P-values (*p*), Cohen’s *d* effect size (*cD*) and *AU C* values are reported using Kolmogorov–Smirnov distance (*d*_*KS*_) and match distance (*d*_*M*_) for the three evaluation measurements computed. Mean and standard deviations of intraclass (in real cases) and interclass distance values are also displayed on the right.

| | | $p$ | $cD$ | $AUC$ | *real vs real* | *real vs phantom* |
| --- | --- | --- | --- | --- | --- | --- |
| Intensity | $d_{KS}$ | 0.172 | 0.267 | 0.607 | 0.024±0.015 | 0.020±0.015 |
| | $d_M$ | 0.706 | 0.049 | 0.530 | 0.224±0.148 | 0.231±0.144 |
| Dispersion | $d_{KS}$ | 0.458 | 0.190 | 0.558 | 0.073±0.040 | 0.065±0.040 |
| | $d_M$ | 0.305 | 0.252 | 0.581 | 5.749±3.005 | 4.919±3.328 |
| Relation<br>art–air | $d_{KS}$ | 0.346 | 0.241 | 0.574 | 0.215±0.135 | 0.245±0.122 |
| | $d_M$ | 0.271 | 0.358 | 0.586 | 8.489±5.140 | 11.051±7.324 |

## 4. Discussion

Both qualitative and quantitative results suggest high similarity between the real and synthetically generated lungs. Therefore, despite the artificial structure of air and blood flow systems in synthetic images, their intensity distributions, dispersions of structures, and relationships between arteries and airways are close enough to real cases. The simulation of features displayed in real CT images of the lungs, such as intertwined structures, partial volume effects affecting small elements, and the limitation in the distinction of medium-to-small generations of the bronchial tree, increases the degree of realism in the final phantoms. Furthermore, the variability introduced by the random component of the system guarantees the ability to generate different and suitable phantoms sharing the same input parameters.

All these characteristics support the capability of the proposed framework to generate databases of labeled anthropomorphic CT images of the lung that could be useful to train and/or evaluate algorithms for processing pulmonary CT images. For example, registration methods could demonstrate their robustness when only internal pulmonary structures have changed. Validation schemes for vessel segmentation algorithms could increase their accuracy and evaluation power with the inclusion of information about the whole structure of interest and its inherent topology. Additionally, differentiation between arteries and veins in the synthetic lungs could encourage and facilitate the development of AV segmentation algorithms, a very promising field with vast potential scientific opportunities, but barely explored until now [36, 37, 38, 16, 39].

On the other hand, the system presents several limitations. Real pulmonary vasculature and airways have a strict tree structure in which each air and blood flow system is connected through the hilum and develops from there. However, lung segmentations exclude this anatomical area, so segmented vessels and airways in the region of interest of real cases are not necessarily connected with the remaining subtrees belonging to the same internal structure. The proposed anthropomorphic phantoms are generated conforming a single connected component, so the spatial distributions of first generations could be far from the real situation. Similarly, the segmental property of the lung is not fully addressed in this work. Different lobes and segments represent nearly independent functional and anatomical units, and they are associated with one principal artery and its accompanying airway. Although real locations of arteries are used in our simulation of the synthetic ones, and oxygenation maps encourage the growth of the structure close to the real one, the random components of the system do not force the generation of one principal artery in each lung segment or lobe. Both situations could be overcome in the future with the inclusion of more complete lung segmentations. A labeled hilum could set the area where all connected air and blood flow systems could start growing. Lobe segmentation could be used to generate subtrees independently in each lobar region.

Apart from these challenges, differences between air and blood flow systems in some features such as branching angles, and the lengths and sizes of segments within the tree have not been studied in depth when generating the phantoms. Although many works [40, 41] have examined these kinds of morphometric parameters, variations and differences in healthy and pathological cases are still being studied using medical imaging, especially in arterial and venous flow systems independently. The development of AV segmentation algorithms would allow the study of morphological and topological properties of pulmonary vessels that could enable the proposed methods to achieve more realistic pulmonary phantoms.

The use of more sophisticated or dedicated methods for pulmonary vessel/airway generation, integrated with the methodology presented in this work to combine different air and blood flow systems could also improve the outcomes. For instance, approaches based on fractal branching algorithms seem to create more realistic pulmonary structures. However, they are less flexible in being able to include some desired pathological variations, which is a critical feature to create a diverse database.

Additionally, it is known that many types of cardiopulmonary pathology present important phenotypical changes associated with structural and image-based features. For example, pulmonary arterial hypertension and bronchiectasis result in abnormal sizes in some of the structures and probably also in branching patterns and general topology [43]. The proposed framework is flexible enough to simulate these states by incorporating the correct information through the input parameters. Other diseases, such as pulmonary embolism and emphysema, could also be simulated in the stage of inclusion of realistic features by incorporating a specific model of the phenotypical imaging expression of the disease. Therefore, despite the described limitations of the system, the ability to tune parameters easily in generating pathological databases increases its utility and power.

The distribution of the labeled synthetic database will enable the community to improve evaluation schemes, to train or test different methods for processing pulmonary structures in CT images, as well as support the development of new image processing algorithms.

## Supporting Information

### S1 Video

Slice-by-slice fly-through of an anthropomorphic pulmonary CT phantom. The resulting CT image and the same image with an accompanying overlay of the synthetic inner structures are shown.

## Acknowledgments

We would like to thank Eduardo Fraile and Patricia Fraga at Unidad Central de Radiodiagnóstico (Madrid, Spain) for assistance during the AV manual labeling; German Gonzalez and George R. Washko at Surgical Planning Laboratory, Brigham and Women’s Hospital (Boston, MA, USA) for supporting the development of the project, and for providing access to the CT cases and their corresponding segmentations, respectively.

## Author Contributions

Conceived and designed the experiments: DJC RSJE MJLC. Performed the experiments: DJC. Analyzed the data: DJC. Contributed reagents/materials/analysis tools: DJC MDC. Wrote the paper: DJC RSJE MJLC.

## Data Availability

CT cases used in this work are part of COPDGene study and can be obtained through dbGap (http://www.ncbi.nlm.nih.gov/projects/gap/cgi-bin/study.cgi?study_id=phs000179.v4.p2).

The whole dataset of anthropomorphic pulmonary CT phantoms (24 synthetic cases) is available via Zenodo at http://dx.doi.org/10.5281/zenodo.20766 (doi:10.5281/zenodo.20766).

